# The genome of the charismatic sea star *Linckia laevigata*

**DOI:** 10.1101/2025.06.09.658723

**Authors:** Caleb J Trimble, Christopher E Laumer, Miles Lamare, Hugh F Carter, Maria Byrne, Matthias Fellner, Suzanne T Williams, Nathan J Kenny

## Abstract

*Linckia laevigata* is a tropical sea star commonly found throughout the Indian and Pacific Oceans, and is one of the top ten-most collected invertebrates, often encountered in the aquarium trade. It has been the subject of investigations into population structure, biodiversity and ecology, particularly regarding gene flow among populations throughout its range, the status of different colour morphs and its relationship with its putative sister species, *L. multifora*. Here we present and describe a high-quality genome assembly for *L. laevigata.* Our assembly is 585.97 Mb in length, with a scaffold N50 of 3 Mb. The genome has a typical repeat (36.99%) and GC content (41.33%), when compared with other echinoderm datasets. Our genome annotation recovers 16,178 genes, with high (89.4%) recovery of the metazoan BUSCO set. This novel resource will provide a model organism for studying the biogeography of the tropical Indo-West Pacific region, and more specifically facilitate the investigation of a range of sea star traits at the genomic level.

**Significance:** Sea stars remain poorly characterised at a genomic level. Data from the charismatic tropical sea star *Linckia laevigata* will enable future studies into the biology, population structure and evolution of these ecologically important species.

## Introduction

The sea star *Linckia laevigata* (Fig. 1a) is a common and charismatic member of the Indo-Pacific coral reef fauna, notable for its striking and variable colouration. A range of molecular studies examining the population structure of the species across its tropical Indo-West Pacific range (Fig. 1b) have used allozyme electrophoresis, analysis of restriction fragment length polymorphisms (RFLP) of the mitochondrial control region, or cytochrome oxidase I (*cox1*) sequence data (e.g. (Williams and Benzie 1998 Juinio-Meñez et al. 2003, Kochzius et al. 2009, Crandall et al. 2008, 2014). Despite these efforts, investigations into the molecular physiology, biogeography and population structure of this species has been hampered by a paucity of molecular data.

**Fig. 1.**
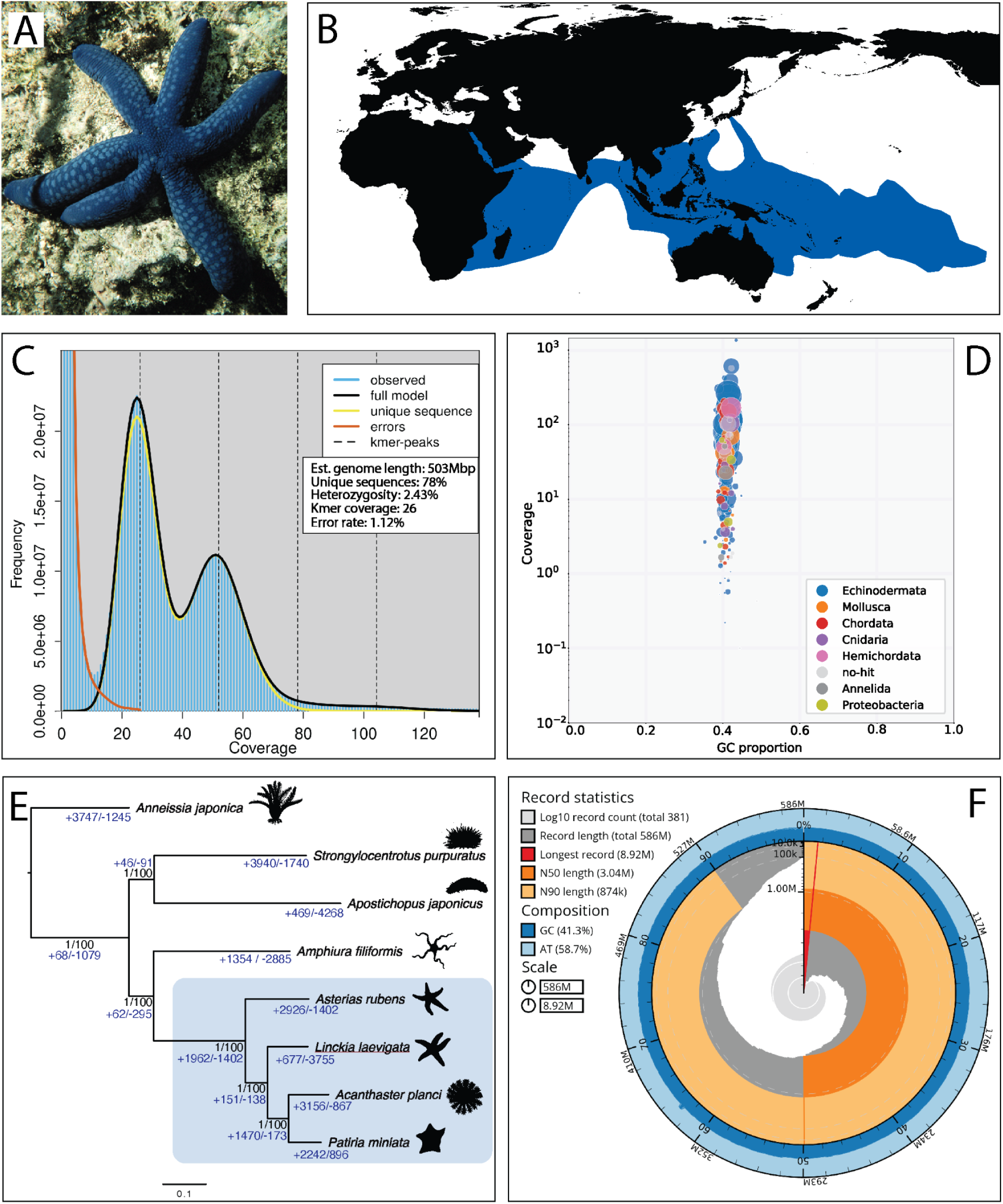
a) *Linckia laevigata,* imaged in the shallow waters of the Great Barrier Reef (Photo: Suzanne Williams). b) Map of the Indo West-Pacific region with areas where *L. laevigata* has been recorded shaded blue (from Williams et al. 2025). c) GenomeScope2.0 (Ranallo-Benavidez et al. 2020) fitted model *k* mer spectra. d) BlobTools2 BlobPlot quantifying coverage and GC content, coloured by sequence similarity binned to phylum e) Species tree with maximum likelihood bootstrap and Bayesian posterior probability support labelled in black text on tree nodes. Values for orthogroup loss and gain per-branching event and per-species are labelled in blue throughout the tree. Species images obtained from www.phylopic.org, image authors noted in Acknowledgements. Images are for illustrative purposes only and do not depict the actual species mentioned. f) BlobTools2 Snail Plot displaying *L. laevigata* genome assembly statistics.

To date, published molecular resources for the *Linckia* genus, beyond the small number of loci used in population studies, are limited to mitochondrial genomes (Hiruta et al. 2020, Inoue et al. 2020) and microsatellites (Yasuda et al. 2012). For a small number of other sea star species, genome sequences are available, including those of *Acanthaster planci* (Hall et al. 2017), *Patiriella regularis* (Long et al. 2016), *Astropecten irregularis* (Adkins et al. 2024), *Luidia sarsii* (Mrowicki et al. 2023), *Marthasterias glacialis* (Lawniczak 2021), *Patiria miniata* (Telmer et al. 2024), *Patiria pectinifera* (Rhee et al. 2024), and *Asterias rubens* (NCBI Bioproject: PRJEB33974). These have been useful for gaining a general understanding of the genetic makeup of sea stars, alongside classical cytogenetic means of investigating genomic content (e.g. Saotome & Komatsu, 2002, Kondo & Akasaka 2012). There are, however, no genomic or transcriptomic resources for members of the family Ophidiasteridae, including *L. laevigata*. Availability of such an assembly could benefit investigations into genetic diversity of sea stars, enabling the isolation of suitable population genetic markers for further study of this iconic species, and improving our understanding of the group more generally. Here, we present a draft genome sequence of *L. laevigata*, as well as gene and repeat annotation, and associated assembly statistics.

## Results and Discussion

### Pre-processing and genome size estimation

DNA was extracted from a single specimen of *L. laevigata* of unknown sex, using the Flow Cell Wash kit to maximize outputs. Circa 37 Gbp of QC-passing LSK114 nanopore reads of 15-20 kbp were obtained from three GridION flow cells.

Jellyfish (Marcais & Kingsford 2011) provided a 21-mers distribution count of *k*-mers within our reads, which exhibits two peaks in the 20x coverage and 40x coverage regions, indicating the heterozygous (2.43%) and diploid nature of the *L. laevigata* genome (Fig. 1c). Sequencing yielded 26-fold haploid *k*-mer coverage on average, and 1.12% of bases as erroneous, consilient with the “Q20” nanopore chemistry used.

The genome has a calculated assembly size of 585.97 Mb, with 381 contigs, 373 of which are larger than 50,000 bp. BUSCO reporting of the *L. laevigata* genome assembly reveals 99.27% recovery of full and partial core metazoan genes. The BUSCO report suggests a highly complete *L. laevigata* genome containing nearly all metazoan core genes, in line with other recently published asteroid genomes.

BlobToolKit and BlobTools2 (Challis et al. 2020) generated a BlobPlot visualising the sequence content of the assembled *L. laevigata* genome (Fig. 1d). Sequences were plotted based on fold coverage and GC proportion, then coloured by phylum showing the closest sequence similarity. The predominant phylum (65.4%) matched in a Diamond BLASTx search versus Uniprot (Buchfink et al. 2015) was Echinodermata, as expected. The majority of data points plotted near 40% GC content on the x-axis with a genomic GC content of 41.33%. Genome GC content near 40% is typical of asteroid genomes; *Acanthaster planci* at 41.31% (Hall et al. 2017), *Astropecten irregularis* at 40.1% (Adkins et al. 2024), *Luidia sarsii* at 44.3% (Mrowicki et al. 2023), *Marthasterias glacialis* at 39.5% (Lawniczak 2021), *Patiria pectinifera* at 39.9% (Rhee et al. 2024), and *Patiria miniata* at 40.5% (Telmer et al. 2024). While there is variation in the coverage of the plotted data in the BlobPlot, the larger quantity of echinoderm datapoints sit around/above 10^2^ coverage.

In the species tree (Fig. 1e) we show the phylogenetic grouping of several species of phylum Echinodermata. *Linckia laevigata* was recovered in a clade of sea stars, as sister to *Patiria miniata* and *Acanthaster planci*. The tree structure is concurrent with systematic relationships present in recent literature (Cohen et al. 2003, Arizza et al. 2017), and has strong statistical support from Bayesian (1,000,000 generations) and Maximum Likelihood (1,000 bootstrap replicate) analyses. The Orthofinder (Emms & Kelly 2019) duplications per-species-tree-node output can be found in Supplementary Table 6.

### Genome assembly statistics

**Table 1.** General statistics on the *Linckia laevigata* genome assembly and annotation. BUSCO score key: C = Complete, S = Single sequence match, D = Duplicated sequence match, F = Fragmented sequence, M = Missing sequence.

| Statistic | Result |
| --- | --- |
| Total Length | 585,975,304 bp |
| Number of Input Reads | 3,332,931 |
| Number of Contigs | 381 |
| N50 | 3,037,324 |
| L50 | 62 |
| GC content (%) | 41.33 |
| Repeat Elements (%) | 36.99 |
| # Genes | 16,178 |
| # mRNAs | 18,972 |
| BUSCO scores *Linckia* genome assembly | C:98.7%[S:97.5%,D:1.2%],F:0.6%,M:0.7% |
| BUSCO scores *Linckia* gene annotation | C:89.4%[S:78.3%,D:11.1%],F:2.0%,M:8.6% |

The total repeat content of the *L. laevigata* genome was 36.99% or ∼216 Mb. The most predominant TE repeats are TIR Elements, consisting of TE species CACTA (4.62%), Mutator (6.66%) hAT (3.46%) and PIF_Harbinger (1.54%). Helitron repeats represent the single largest class of TE repeats within the *L. laevigata* genome at 6.53%. The repeat content of the *L. laevigata* genome is within the typical range for eukaryotic genomes as well as echinoderm genomes (Liao et al. 2023) (See Supplementary Table 1 and Supplementary Figure 1 for more information). Interestingly, no SINE elements from current databases were retrieved in the *L. laevigata* genome, save for a small quantity of tRNA-derived sequences.

### Gene prediction

BRAKER3 gene prediction recovered 16,178 genes and 18,972 mRNAs, with an average of 1.2 mRNAs per gene in the *L. laevigata* genome, a slightly smaller number than that seen in most other echinoderm genomes to date. BUSCO reporting for transcript sequences yielded 89% total recovery of core metazoan genes, with 78% single copy, 11% multiple-copy, and 2.0% fragmented genes. We suspect that this smaller gene count might be due to incomplete recovery of novel and lineage-specific gene sequences due to a lack of reference Ophidiasteridae sequences for annotation. However, BUSCO reporting suggests a high quality geneset containing most but not all metazoan core genes, which gives us confidence in our annotation of genes shared more widely across the animal tree of life.

### Functional annotation

Gene Ontology (GO) terms were mapped to 56.35% of annotated sequences in the *L. laevigata* genome using OmicsBox (OmicsBox, 2019), with a total of 197,992 annotations and an average of 18.52 GO terms per sequence. Functional categorization noted 15.04% of sequences involved in information storage and processing, 30.34% in cellular processes and signaling, and 18.53% in metabolism. Poorly characterised sequences with ‘unknown’ functions made up the remaining 25% of gene models. This is expected for *L. laevigata*, as echinoderms are not well-represented in extant databases, resulting in gaps in annotation for some gene families. Orthologous group distribution revealed Eukaryota (18.52%), Opisthokonta (16.86%), and Metazoa (16.7%) with minimal representation in bacteria (0.53%). The EggNOG (Huerta-Cepas et al. 2019) orthologous group distribution for *L. laevigata* is representative, reflecting the evolutionary placement of the sea star within class Asteroidea.

### Gene loss and gain statistics

Gene gain and loss across the phylogeny containing the seven species of echinoderm shown in Figure 1e were calculated using CAFE5 (Mendes et al. 2020). *Linckia laevigata* shows a relatively high gene loss value, and lowered gene gain value in comparison to other echinoderm species. This could be partly due to the genome annotation process, which, as noted above, may have been conservative in its assignment of genes to annotations, and some novel genes may have been missed.

The genome of sea cucumber *Apostichopus japonicus* was the only species to show greater gene loss than *L. laevigata*.

### General Discussion and Conclusions

The class Asteroidea comprises a diverse range of globally distributed species, many of which are ecologically influential (e.g. Matthews et al. 2024). These keystone species have driven significant biogeographic change, as seen by the decimation of coral by population outbreaks of *Acanthaster* spp. (Matthews et al. 2024). These ecological impacts underscore the importance of the Asteroidea for ecological and evolutionary research, and the fundamental information provided by genomic resources is vital for delving into a range of scientific questions. The resource presented here will be a valuable addition to the published record.

Genomes also provide important data for the study of systematic relationships of groups. The systematics of Asteroidea is not fully resolved, with understanding of the phylogenetic relationships of some groups impeded by paucity of sufficient sequencing data (Linchangco et al. 2017). Recent studies have used mitochondrial genomes and transcriptomic resources to rectify this, but although these are useful, they do not fully resolve relationships, and many currently recognised families and orders are recovered as polyphyletic (Mah and Foltz 2011, Jossart et al. 2024). Combining previous molecular data with new genomic information will contribute to improved systematics within Asteroidea. Our genome is the first to be sequenced for ophidiasterid sea stars, which are already often represented by *L. laevigata* in molecular phylogenetic studies (e.g. Jossart et al. 2024), rendering this genomic resource ideal as a basis for future work.

Furthermore, *L. laevigata* are also of interest in and of themselves, as they are highly desirable in the aquarium and curio trades, and are heavily fished at some localities (Wabnitz 2003); *L. laevigata* has been reported as one of the top ten-most collected invertebrates. This could lead to excessive exploitation, and genomic resources will be useful to inform conservation studies, to track and manage population numbers and anthropogenic impacts. This genome will therefore be of practical utility for this species into the future.

## Materials and Methods

### Sampling and lab work

A skin biopsy (as described in Williams et al. 2025) was taken from a specimen collected from Lizard Island, Queensland, Australia (Permit G14/36625.2). Igh molecular weight genomic DNA was extracted using the Monarch® HMW DNA Extraction Kit for Tissue (New England Biolabs cat. no. T3060S), as per manufacturer’s instructions. Quality controls using a dsDNA Qubit (ThermoFisher) and Genomic DNA TapeStation (Agilent) assay indicated the presence of ≥5 μg of high molecular weight DNA (>50% above 40 kb). Genomic DNA was sheared using 20 passes through a 27 G needle and loaded onto a BluePippin BLF7510 cassette for 10 kb size selection, followed by NEBNext FFPE and end-repair reactions and nanopore adapter ligation with the LSK114 ligation kit, as per manufacturer’s recommendations. DNA was sequenced across 3 GridION R10.4.1 flow cells with SUP base calling, loading 20 fmol per run, and employing the flow cell wash kit (EXP-WSH004) to reload two flow cells for a total of 5 library loadings, generating 25 GB of raw data with a read N50 of 20 kb. A second, more highly-sheared library was also generated and loaded on the same flow cells, generating a further 18.25 GB of N50 15 kb data (Supplementary Table 2).

### Genome assembly

Reads passing the MinKNOW base call threshold (Q10) were adapter trimmed using Porechop_abi (Bonenfant et al. 2023). These were used as input for assembly in NextDenovo (Hu et al. 2024), a “correct then assemble” string graph assembler that has been shown to perform well for highly heterozygous genomes with nanopore data (Sun et al. 2021). Following assembly, we applied NextPolish to improve the base consensus quality, mapping the same reads used in assembly, and following the default configuration files provided in the documentation. Finally, the adapter-trimmed reads were again mapped to the NextPolish finished assembly using minimap2 (-x map-ont), and the manual purge_dups pipeline (Guan et al. 2020) was utilized to purge any remaining haplotigs from the NextDenovo assembly (218 contigs, spanning 28.89 Mb of data).

Genome size estimates were attained by measuring average coverage of the assembly, and by counting *k* mers within our reads using Jellyfish v2.3.1 (Marcais et al. 2011) before running GenomeScope 2.0 (Ranallo-Benavidez et al. 2020) online. The resulting assembly was assessed using BUSCO v5.7.1 (Manni et al. 2021) on euk_gen and euk_tran modes, utilising the Metazoa_odb10 database containing 954 BUSCOs (Supplementary Table 3). Dependencies were hmmsearch 3.1, metaeuk: 6.a5d39d9.

Repeat masking was performed using EDTA (Ou et al. 2019) version 2.1.3 on sensitive mode and RepeatMasker version 4.1.1 (Smit et al. 2013) . EDTA identifies and quantifies repeat regions within raw read data (see more at https://github.com/oushujun/EDTA). From EDTA and RepeatMasker we obtained a curated repeat library annotated for species of Transposable Elements (TE), as well as a repeat-masked output file for later use in downstream analyses.

The gene content of the *L. laevigata* genome was predicted using the Braker3 v3.0.3 annotation pipeline (Hoff et al. 2016, Stanke et al. 2008). The *Asterias rubens* proteome was sourced from NCBI (https://www.ncbi.nlm.nih.gov/genome/annotation_euk/Asterias_rubens/100/) to be used as a training dataset. *Linckia laevigata* Illumina transcriptomes and a *L. laevigata* MAS-Seq transcriptome (Laumer, supplied) were used for evidence during annotation; mapped to the genome assembly with minimap v2.26-r1175 (Li 2018). Functional annotation of the Braker3 predicted gene set was generated using the OmicsBox graphical user interface (OmicsBox, 2019). EggNOG (Huerta-Cepas et al. 2019) analysis was carried out to determine gene ontology (GO) and orthology.

Statistics for the genome assembly were obtained using QUAST (Gurevich et al. 2013, Supplementary Table 4), and agat_sp_functional_statistics.pl from the AGAT package (Dainat 2024, Supplementary Table 5) was used to generate statistics for the genome annotation.

The BlobPlot (Challis et al. 2020) (Fig. 1d) was generated using the Diamond package (Buchfink et al. 2015) and BLASTx (Camacho et al. 2009) to align the *L. laevigata* genome assembly against the NIH Non-Redundant Standard Protein database (Downloaded August 2024).

Our species tree (Fig. 1e) was constructed using OrthoFinder (Emms & Kelly 2019) -derived single copy orthogroups and inferred with maximum likelihood in IQ-TREE (Minh et al. 2020) under the LG+I+F model with 1,000 bootstrap replicates. Bayesian support was also provided, using MrBayes (Ronquist et al. 2012) with 1,000,000 generations, a sampling rate of 100, and a burn-in of 2,500. Besides our gene annotations from *L. laevigata*, six additional echinoderm species with reference genomes on public databases (NCBI and Echinobase) were selected to be used in generating the species tree: *Strongylocentrotus purpuratus, Apostichopus japonicus, Amphiuria filiformis, Asterias rubens, Acanthaster planci,* and *Patiria miniata*. The crinoid feather star *Anneissia japonica* was used as the outgroup based on previous studies (Cohen et al. 2003, Arizza et al. 2017). Input files were all available reference quality echinoderm genome translated protein fasta files downloaded from the NCBI and Echinobase databases. Cafe5 (Mendes et al. 2020) was used to calculate values for gene loss and gain between echinoderm species from the Orthofinder output data. The Orthofinder SpeciesTree_rooted.txt output was converted to an ultrametric format with Orthofinder’s make_ultrametric.py script. Additionally, the Orthofinder Orthogroups.GeneCount.tsv output was edited with the cafetutorial_clade_and_size_filter.py script, and these files were used as inputs for Cafe5 analysis. Cafe5 was run with default settings, with the resulting gene gain/loss values incorporated into Figure 1e.

## Acknowledgements

We acknowledge the Traditional Owners of the area from which our samples were taken, the Dingaal people. We pay our respects to their Elders past, present, and emerging, and acknowledge their ongoing connection to Country. We thank Andrea Waeschenbach for providing the tissue sample of *Linckia laevigata* and Anne Hoggett and Lyle Vail for fieldwork assistance on Lizard Island. Catalyst: Seeding funding for this work (23-UOO-003-CSG to NJK (PI), STW, MB and ML) was provided by the New Zealand Ministry of Business, Innovation and Employment and administered by the Royal Society Te Apārangi. Chris Laumer is funded by a Royal Society University Research Fellowship, award no. URF\R1\221744. This work was part-funded by a grant from the National Geographic Society (https://nationalgeographic.org) to STW (NGS-52123R-19) and NHM Science Investment Funds to STW. Sampling on Lizard Island was covered under permit G19/39553.1 (held by Lizard Island Research Station of the Australian Museum). We thank the authors of the images used in Fig 1e: *Acanthaster planci* CC0 1.0 Zechariah Meunier, Asteriidae CC BY 3.0 Mali’o Kodis based on photograph by "User: Wildcat Dunny" (http://www.flickr.com/people/wildcat_dunny/), “*Holothuria*” CC0 1.0 Lauren Sumner-Rooney; *Patiria pectinifera* CC0 1.0 Kanchi Nanjo; Ophiuroidea CC0 1.0 Kurtis Wothe; *Arbacia punctulata* CC BY 4.0 Julia Notar; *Asterias rubens* CC BY-SA 3.0 by Hans Hillewaert (photo) and T. Michael Keesey; and *Antedon bifida* CC0 1.0 by Guillaume Dera.

## Author Contributions

This study was conceived by STW and NJK with contributions from CEL. CEL undertook sequencing and assembly tasks, and CJT performed annotation and further analysis of genome content. All authors contributed to analysis of results and writing of text. The final version of the manuscript was approved by all authors.

## Data Availability

Transcriptome raw reads are available from the NCBI SRA under SRX2728034. Genome raw reads are available from the NCBI SRA under Bioproject PRJNA1269645. Genome assembly and related files available from https://data.nhm.ac.uk/dataset/edit/genomic-resources-for-linckia-laevigata.

## Supplementary Material

**Supplementary Table 1.** EDTA tabular output detailing the transposable element content of the *Linckia laevigata* genome. Input file *Linckia laevigata* genome assembly fasta.

| TE Class | Number of Elements | Length (bp) | Percentage of Sequence |
| --- | --- | --- | --- |
| DNA transposon | 20 | 15348 | 0.00% |
| LINE | -- | -- | -- |
| CR1 | 5979 | 1971212 | 0.34% |
| I | 48 | 47856 | 0.01% |
| Jockey | 379 | 168443 | 0.03% |
| L1 | 1706 | 1435632 | 0.24% |
| L2 | 8484 | 3207951 | 0.55% |
| Proto2 | 106 | 53891 | 0.01% |
| R2 | 233 | 143317 | 0.02% |
| R4 | 649 | 248078 | 0.04% |
| RTE | 3774 | 1649843 | 0.28% |
| Rex | 1691 | 752240 | 0.13% |
| unknown | 1263 | 456627 | 0.08% |
| LTR | -- | -- | -- |
| Bel_Pao | 261 | 115528 | 0.02% |
| Copia | 250 | 92671 | 0.02% |
| Gypsy | 20296 | 14283631 | 2.44% |
| unknown | 77816 | 26955388 | 4.60% |
| SINE | -- | -- | -- |
| tRNA | 6 | 504 | 0.00% |
| TIR | -- | -- | -- |
| CACTA | 92156 | 27057372 | 4.62% |
| Mutator | 123975 | 39010812 | 6.66% |
| P | 27 | 13289 | 0.00% |
| PIF_Harbinger | 33086 | 8997636 | 1.54% |
| Tc1_Mariner | 8329 | 1325994 | 0.23% |
| Transib | 202 | 71138 | 0.01% |
| hAT | 63840 | 20246107 | 3.46% |
| piggyBac | 29 | 28861 | 0.00% |
| polinton | 1351 | 1918991 | 0.33% |
| Low complexity | 439 | 106964 | 0.02% |
| nonLTR | -- | -- | -- |
| DIRS_YR | 795 | 663740 | 0.11% |
| Ngaro_YR | 247 | 142570 | 0.02% |
| Penelope | 4047 | 1743798 | 0.30% |
| nonTIR | -- | -- | -- |
| helitron | 149480 | 38248172 | 6.53% |
| Repeat fragment | 77844 | 25584818 | 4.37% |
| total interspersed | 678808 | 216758422 | 36.99% |
| Total | 678808 | 216758422 | 36.99% |
| Total Sequences: | 381 |  |  |
| Total Length: | 585975304 bp |  |  |

**Supplementary Table 2.**
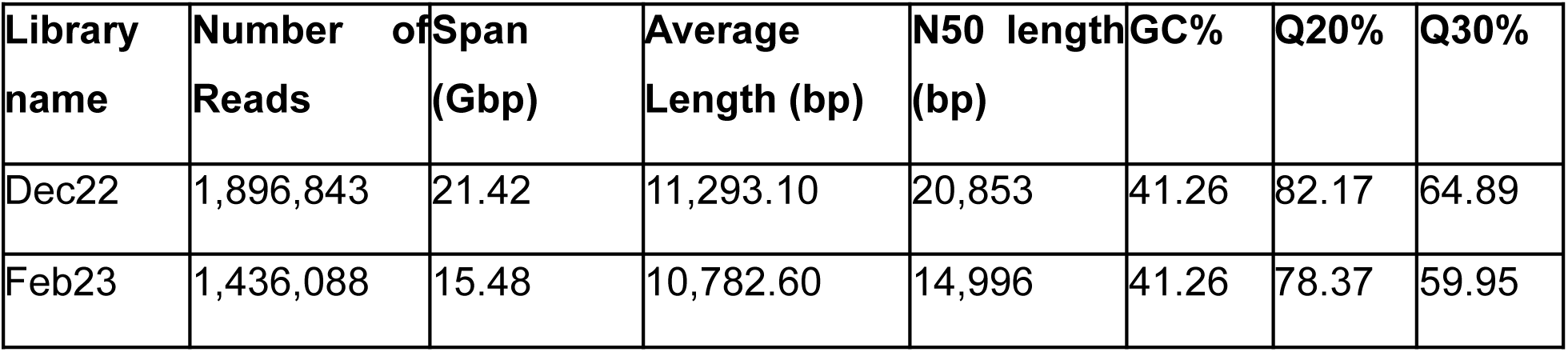
Summary statistics describing long-read data outputs from two libraries prepared using the same genomic DNA extraction.

| Library name | Number of Reads | Span (Gbp) | Average Length (bp) | N50 length (bp) | GC% | Q20% | Q30% |
| --- | --- | --- | --- | --- | --- | --- | --- |
| Dec22 | 1,896,843 | 21.42 | 11,293.10 | 20,853 | 41.26 | 82.17 | 64.89 |
| Feb23 | 1,436,088 | 15.48 | 10,782.60 | 14,996 | 41.26 | 78.37 | 59.95 |

**Supplementary Table 3.**
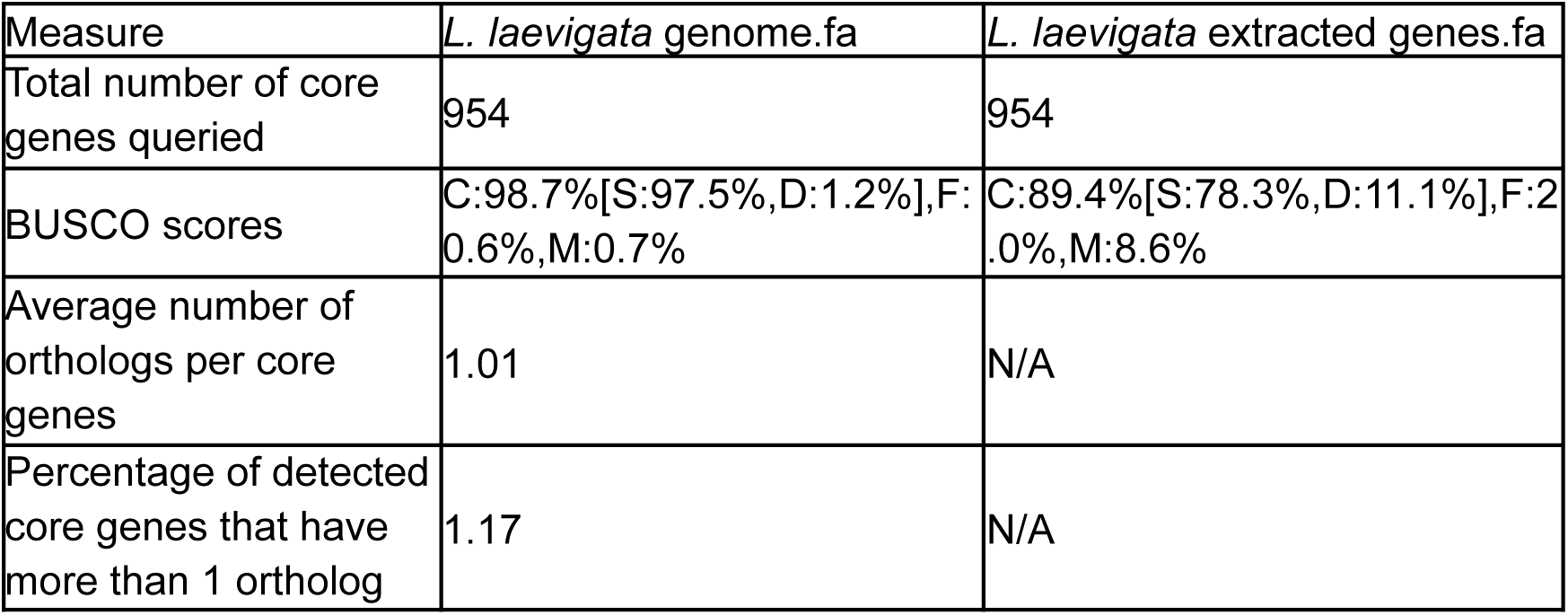
Table of BUSCO scores and statistics using the *Linckia laevigata* genome assembly and extracted Braker3 geneset. BUSCO score key: C = Complete, S = Single sequence match, D = Duplicated sequence match, F = Fragmented sequence, M = Missing sequence.

| Measure | *L. laevigata* genome.fa | *L. laevigata* extracted genes.fa |
| --- | --- | --- |
| Total number of core genes queried | 954 | 954 |
| BUSCO scores | C:98.7%[S:97.5%,D:1.2%],F:0.6%,M:0.7% | C:89.4%[S:78.3%,D:11.1%],F:2.0%,M:8.6% |
| Average number of orthologs per core genes | 1.01 | N/A |
| Percentage of detected core genes that have more than 1 ortholog | 1.17 | N/A |

**Supplementary Table 4.** Quast tabular output detailing contig and assembly statistics of the *L. laevigata* genome. All statistics are based on contigs of size >= 500 bp, unless otherwise noted (e.g., "# contigs (>= 0 bp)" and "Total length (>= 0 bp)" include all contigs).

| Assembly | *Linckia laevigata* assembly |
| --- | --- |
| # contigs ($\geq 0$ bp) | 381 |
| # contigs ($\geq 1000$ bp) | 381 |
| # contigs ($\geq 5000$ bp) | 381 |
| # contigs ($\geq 10000$ bp) | 381 |
| # contigs ($\geq 25000$ bp) | 380 |
| # contigs ($\geq 50000$ bp) | 373 |
| Total length ($\geq 0$ bp) | 585,975,304 |
| Total length ($\geq 1000$ bp) | 585,975,304 |
| Total length ($\geq 5000$ bp) | 585,975,304 |
| Total length ($\geq 10000$ bp) | 585,975,304 |
| Total length ($\geq 25000$ bp) | 585,956,419 |
| Total length ($\geq 50000$ bp) | 585,696,600 |
| # contigs | 381 |
| Largest contig | 8,918,583 |
| Total length | 585,975,304 |
| GC (%) | 41.33 |
| N50 | 3,037,324 |
| N90 | 874,130 |
| auN | 3,392,528.2 |
| L50 | 62 |
| L90 | 196 |
| # N's per 100 kbp | 0 |

**Supplementary Table 5.** Output table from the AGAT (Dainat J. 2024) bioinformatics package script, agat_sp_functional_statistics.pl, featuring average gene size, exon number, and additional statistics on the *L. laevigata* Braker3 genome annotation.

| Metric | Value |
| --- | --- |
| Number of genes | 16,178 |
| Number of transcripts | 18,972 |
| Number of cds | 18,972 |
| Number of exons | 140,681 |
| Number of introns | 121,709 |
| Number of start_codons | 18,962 |
| Number of stop_codons | 18,964 |
| Number of exons in cds | 140,681 |
| Number of introns in cds | 121,709 |
| Number of introns in exons | 121,709 |
| Number genes overlapping | 82 |
| Number of single exon genes | 3,086 |
| Number of single exon transcripts | 3,362 |
| Mean transcripts per gene | 1.2 |
| Mean cds per transcript | 1 |
| Mean exons per transcript | 7.4 |
| Mean introns per transcript | 6.4 |
| Mean start_codons per transcript | 1 |

**Supplementary Table 6.** Table of duplications per species-tree-node from Orthofinder astral consensus gene tree outputs. Row labels “N0-N6” refer to nodes within the tree.

| Species Tree Node | Duplications (all) | Duplications (50% support) |
| --- | --- | --- |
| *Strongylocentrotus purpuratus* | 20,500 | 20,500 |
| *Asterias rubens* | 8,901 | 8,901 |
| *Amphiura filiformis* | 11,653 | 11,653 |
| *Apostichopus japonicus* | 4,976 | 4,976 |
| *Acanthaster planci* | 13,759 | 13,759 |
| *Anneissia japonica* | 14,691 | 14,691 |
| *Linckia laevigata* | 5,398 | 5,398 |
| *Patiria miniata* | 16,648 | 16,648 |
| N0 | 973 | 281 |
| N1 | 824 | 151 |
| N2 | 237 | 56 |
| N3 | 60 | 60 |
| N4 | 693 | 561 |
| N5 | 448 | 332 |
| N6 | 406 | 406 |

**Supplementary Figure 1.**
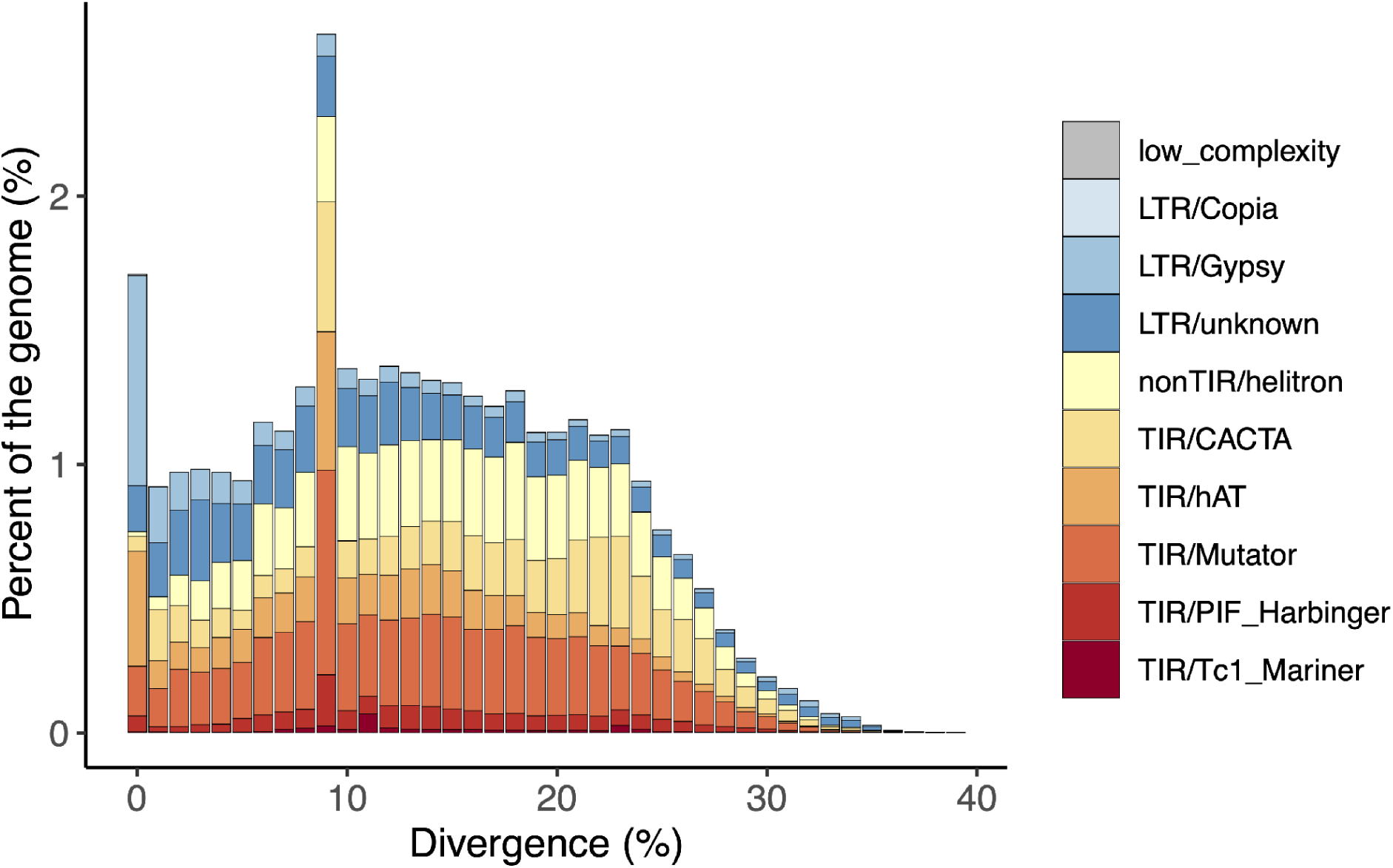
*Linckia laevigata* genome repeat landscape sourced from the EDTA.pl bioinformatics package Ou et al. 2019. TE repeat families are separated by percentage sequence divergence and percentage of genome makeup.

## References

1. Adkins P, Harley J, Bishop J, Marine Biological Association Genome Acquisition Lab, Darwin Tree of Life Barcoding collective, Wellcome Sanger Institute Tree of Life Management, Samples and Laboratory team, et al. The genome sequence of the sand star, *Astropecten irregularis* (Pennant, 1777). Wellcome Open Res. 2024 Aug 7;9:431.

2. Arizza V, Pagliara P, Stabili L, Smith L, Garcia-Arraras J, Henson J, et al. Complexity of the Immune System in Echinoderms. In: Complexity of the Immune System in Echinoderms. Springer Publisher; 2017. p. 1–101.

3. Bonenfant Q, Noé L, Touzet H. Porechop_ABI: discovering unknown adapters in Oxford Nanopore Technology sequencing reads for downstream trimming. Bioinform Adv. 2023;3(1):vbac085.

4. Brůna T, Lomsadze A, Borodovsky M. GeneMark-ETP significantly improves the accuracy of automatic annotation of large eukaryotic genomes. Genome Res. 2024 Jun 25;34(5):757–68.

5. Brůna T, Hoff KJ, Lomsadze A, Stanke M, Borodovsky M. BRAKER2: automatic eukaryotic genome annotation with GeneMark-EP+ and AUGUSTUS supported by a protein database. NAR Genom Bioinform. 2021 Mar;3(1):lqaa108.

6. Buchfink B, Xie C, Huson DH. Fast and sensitive protein alignment using DIAMOND. Nat Methods. 2015 Jan;12(1):59–60.

7. Camacho C, Coulouris G, Avagyan V, Ma N, Papadopoulos J, Bealer K, et al. BLAST+: architecture and applications. BMC Bioinformatics. 2009 Dec 15;10(1):421.

8. Challis R, Richards E, Rajan J, Cochrane G, Blaxter M. BlobToolKit - interactive quality assessment of genome assemblies. G3 (Bethesda). 2020 Apr 9;10(4):1361–74.

9. Cohen BL, Améziane N, Eleaume M, de Forges BR. Crinoid phylogeny: a preliminary analysis (Echinodermata: Crinoidea). Mar Biol. 2004 Mar;144(3):605–17.

10. Crandall ED, Jones ME, Muñoz MM, Akinronbi B, Erdmann MV, Barber PH. Comparative phylogeography of two seastars and their ectosymbionts within the Coral Triangle. Mol Ecol. 2008 Dec;17(24):5276–90.

11. Dainat J, Hereñú D, Murray KD, Davis E, Ugrin I, Crouch K, et al. NBISweden/AGAT: AGAT-v1.4.1. Zenodo; 2024.

12. Emms DM, Kelly S. OrthoFinder: phylogenetic orthology inference for comparative genomics. Genome Biol. 2019 Nov 14;20(1):238.

13. Gotoh O. A space-efficient and accurate method for mapping and aligning cDNA sequences onto genomic sequence. Nucleic Acids Res. 2008 May;36(8):2630–8.

14. Guan D, McCarthy SA, Wood J, Howe K, Wang Y, Durbin R. Identifying and removing haplotypic duplication in primary genome assemblies. Bioinformatics. 2020 May 1;36(9):2896–8.

15. Gurevich A, Saveliev V, Vyahhi N, Tesler G. QUAST: quality assessment tool for genome assemblies. Bioinformatics. 2013 Apr 15;29(8):1072–5.

16. Hall MR, Kocot KM, Baughman KW, Fernandez-Valverde SL, Gauthier MEA, Hatleberg WL, et al. The crown-of-thorns starfish genome as a guide for biocontrol of this coral reef pest. Nature. 2017;544(7649):231-4.

17. Hoff KJ, Lomsadze A, Borodovsky M, Stanke M. Whole-genome annotation with BRAKER. Methods Mol Biol. 2019;1962:65–95.

18. Hoff KJ, Lange S, Lomsadze A, Borodovsky M, Stanke M. BRAKER1: Unsupervised RNA-Seq-based genome annotation with GeneMark-ET and AUGUSTUS. Bioinformatics. 2016 Mar 1;32(5):767–9.

19. Hu J, Wang Z, Sun Z, Hu B, Ayoola AO, Liang F, et al. NextDenovo: an efficient error correction and accurate assembly tool for noisy long reads. Genome Biol. 2024 Apr 26;25(1):107.

20. Huerta-Cepas J, Szklarczyk D, Heller D, Hernández-Plaza A, Forslund SK, Cook H, et al. eggNOG 5.0: a hierarchical, functionally and phylogenetically annotated orthology resource based on 5090 organisms and 2502 viruses. Nucleic Acids Res. 2019 Jan 8;47(D1):D309–14.

21. Iwata H, Gotoh O. Benchmarking spliced alignment programs including Spaln2, an extended version of Spaln that incorporates additional species-specific features. Nucleic Acids Res. 2012 Nov 1;40(20):e161.

22. Jossart Q, Kochzius M, Danis B, Moreau CVE, Schön I, Fourdrilis S, et al. Implications of extensive addition of new mitogenomes for sea star phylogenetics and evolution (Echinodermata: Asteroidea). Zoological Journal of the Linnean Society. 2024;202(4).

23. Juinio-Meñez MA, Magsino RM, Ravago-Gotanco R, Yu ET. Genetic structure of *Linckia laevigata* and *Tridacna crocea* populations in the Palawan shelf and shoal reefs. Mar Biol. 2003 Apr;142(4):717–26.

24. Kim D, Paggi JM, Park C, Bennett C, Salzberg SL. Graph-based genome alignment and genotyping with HISAT2 and HISAT-genotype. Nat Biotechnol. 2019 Aug;37(8):907–15.

25. Kochzius M, Seidel C, Hauschild J, Kirchhoff S, Mester P, Meyer-Wachsmuth I, et al. Genetic population structures of the blue starfish *Linckia laevigata* and its gastropod ectoparasite *Thyca crystallina*. Marine Ecology Progress Series. 2009;396:211–9.

26. Kondo M, Akasaka K. Current Status of Echinoderm Genome Analysis - What do we Know? Curr Genomics. 2012 Apr;13(2):134–43.

27. Kovaka S, Zimin AV, Pertea GM, Razaghi R, Salzberg SL, Pertea M. Transcriptome assembly from long-read RNA-seq alignments with StringTie2. Genome Biol. 2019 Dec 16;20(1):278.

28. Lawniczak MKN, Darwin Tree of Life Barcoding collective, Wellcome Sanger Institute Tree of Life programme, Wellcome Sanger Institute Scientific Operations: DNA Pipelines collective, Tree of Life Core Informatics collective, Darwin Tree of Life Consortium. The genome sequence of the spiny starfish, *Marthasterias glacialis* (Linnaeus, 1758). Wellcome Open Res. 2021 Nov 5;6:295.

29. Li H. Minimap2: pairwise alignment for nucleotide sequences. Bioinformatics. 2018 Sep 15;34(18):3094–100.

30. Liao X, Zhu W, Zhou J, Li H, Xu X, Zhang B, et al. Repetitive DNA sequence detection and its role in the human genome. Commun Biol. 2023 Sep 19;6(1):954.

31. Linchangco GV, Foltz DW, Reid R, Williams J, Nodzak C, Kerr AM, et al. The phylogeny of extant starfish (Asteroidea: Echinodermata) including *Xyloplax*, based on comparative transcriptomics. Molecular Phylogenetics and Evolution. 2017;115:161–70.

32. Long KA, Nossa CW, Sewell MA, Putnam NH, Ryan JF. Low coverage sequencing of three echinoderm genomes: the brittle star *Ophionereis fasciata*, the sea star *Patiriella regularis*, and the sea cucumber *Australostichopus mollis*. Gigascience. 2016 May 10;5(1):20.

33. MAH C, FOLTZ D. Molecular phylogeny of the Valvatacea (Asteroidea: Echinodermata). Zoological Journal of the Linnean Society. 2011;161(4):769–88.

34. Marçais G, Kingsford C. A fast, lock-free approach for efficient parallel counting of occurrences of k-mers. Bioinformatics. 2011 Mar 15;27(6):764–70.

35. Manni M, Berkeley MR, Seppey M, Simão FA, Zdobnov EM. BUSCO update: Novel and streamlined workflows along with broader and deeper phylogenetic coverage for scoring of eukaryotic, prokaryotic, and viral genomes. Mol Biol Evol. 2021 Sep 27;38(10):4647–54.

36. Matthews SA, Williamson DH, Beeden R, Emslie MJ, Abom RTM, Beard D, et al. Protecting Great Barrier Reef resilience through effective management of crown-of-thorns starfish outbreaks. PLOS ONE. 2024;19(4):e0298073.

37. Mendes FK, Vanderpool D, Fulton B, Hahn MW. CAFE 5 models variation in evolutionary rates among gene families. Bioinformatics. 2021 Apr 1;36(22–23):5516–8.

38. Minh BQ, Schmidt HA, Chernomor O, Schrempf D, Woodhams MD, von Haeseler A, et al. IQ-TREE 2: New models and efficient methods for phylogenetic inference in the genomic era. Mol Biol Evol. 2020 May 1;37(5):1530–4.

39. O’Leary NA, Cox E, Holmes JB, Anderson WR, Falk R, Hem V, Tsuchiya MTN, Schuler GD, Zhang X, Torcivia J, et al. Exploring and retrieving sequence and metadata for species across the tree of life with NCBI Datasets. Sci Data. 2024 Jul 5;11(1):732. doi: 10.1038/s41597-024-03571-y. PMID: 38969627; PMCID: PMC11226681.

40. OmicsBox – Bioinformatics Made Easy, BioBam Bioinformatics, March 3, 2019. https://www.biobam.com/omicsbox

41. Ou S, Su W, Liao Y, Chougule K, Agda JRA, Hellinga AJ, et al. Benchmarking transposable element annotation methods for creation of a streamlined, comprehensive pipeline. Genome Biol. 2019 Dec 16;20(1):275.

42. Pertea G, Pertea M. GFF utilities: GffRead and GffCompare. F1000Res. 2020 Apr 28;9:304.

43. Ranallo-Benavidez TR, Jaron KS, Schatz MC. GenomeScope 2.0 and Smudgeplot for reference-free profiling of polyploid genomes. Nat Commun. 2020 Mar 18;11(1):1432.

44. Rhee J-S, Nam S-E, Lee SJ, Park H. De Novo Genome Assembly of the Sea Star Patiria pectinifera (Muller & Troschel, 1842) Using Oxford Nanopore Technology and Illumina Platforms. Diversity. 2024;16(2):91.

45. Ronquist F, Teslenko M, van der Mark P, Ayres DL, Darling A, Höhna S, et al. MrBayes 3.2: efficient Bayesian phylogenetic inference and model choice across a large model space. Syst Biol. 2012 May;61(3):539–42.

46. Saotome K, Komatsu M. Chromosomes of Japanese starfishes. Zoolog Sci. 2002 Oct;19(10):1095–103.

47. Smit AFA, Hubley R, Green P. RepeatMasker Open-4.0. 2013–2015. 2015; Available from: https://scholar.google.com/citations?user=SOqGFioAAAAJ&hl=en&oi=sra

48. Stanke M, Schöffmann O, Morgenstern B, Waack S. Gene prediction in eukaryotes with a generalized hidden Markov model that uses hints from external sources. BMC Bioinformatics. 2006 Feb 9;7(1):62.

49. Stanke M, Diekhans M, Baertsch R, Haussler D. Using native and syntenically mapped cDNA alignments to improve de novo gene finding. Bioinformatics. 2008 Mar 1;24(5):637–44.

50. Sun J, Li R, Chen C, Sigwart JD, Kocot KM. Benchmarking Oxford Nanopore read assemblers for high-quality molluscan genomes. Philos Trans R Soc Lond B Biol Sci. 2021 May 24;376(1825):20200160.

51. Telmer CA, Karimi K, Chess MM, Agalakov S, Arshinoff BI, Lotay V, et al. Echinobase: a resource to support the echinoderm research community. Genetics. 2024;227(1). 10.1093/genetics/iyae002

52. Wabnitz, Colette. 2003. From Ocean to Aquarium: The Global Trade in Marine Ornamental Species. UNEP/Earthprint.

53. Wellcome Sanger Institute. Asterias rubens (common starfish) genome assembly, eAstRub1 [Internet]. Cambridge (UK): Wellcome Sanger Institute; 2019 Aug 16 [cited 2023 Sep 13]. Available from: https://www.ncbi.nlm.nih.gov/bioproject/PRJEB33974/

54. Williams ST, Heyworth SM, Kano Y, Roberts NW, Carter HF, Cheney KL. The blue advantage: a novel blue carotenoprotein pigment in the tropical seastar *Linckia laevigata* is an antioxidant defence against extreme environmental stress. Mar Biol. 2025 Feb;172(2): 31.

55. Williams ST, Benzie JAH. EVIDENCE OF A BIOGEOGRAPHIC BREAK BETWEEN POPULATIONS OF A HIGH DISPERSAL STARFISH: CONGRUENT REGIONS WITHIN THE INDO-WEST PACIFIC DEFINED BY COLOR MORPHS, mtDNA, AND ALLOZYME DATA. Evolution. 1998;52(1):87–99.

56. Yasuda N, Taquet C, Nagai S, Wörheide G, Nadaoka K. Development of microsatellite loci in the common reef starfish *Linckia laevigata* and *Linckia multifora*. Ecol Res. 2012 Nov;27(6):1095–7.

