## Supplementary figures and images for "The genome of the charismatic sea star *Linckia laevigata*"

### Supplementary Figure 1

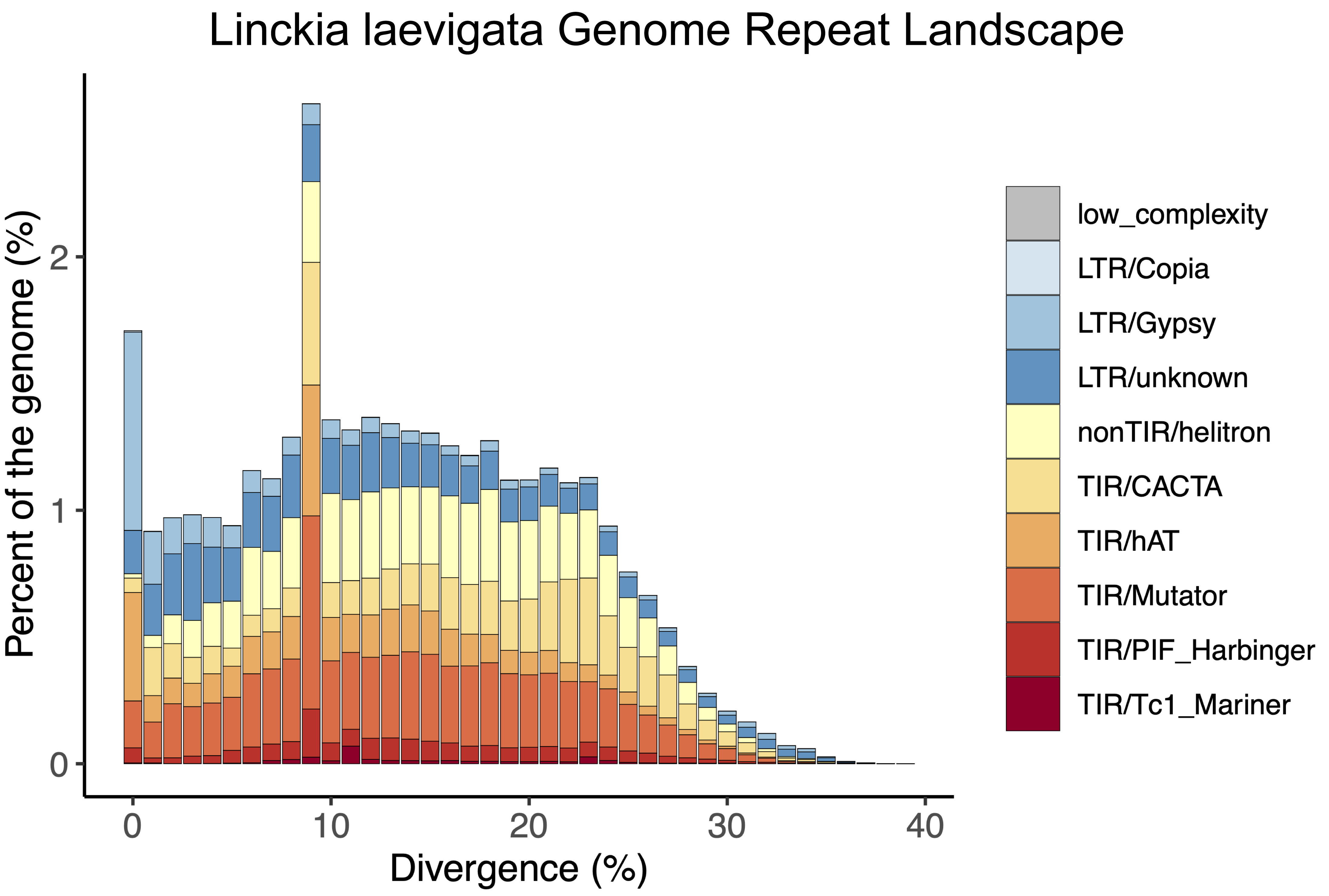
